# Glycerol suppresses lactose-dependent growth of *Streptococcus pyogenes* through a transcription-independent mechanism

**DOI:** 10.1101/2025.11.24.690200

**Authors:** Ayako Bando, Yujiro Hirose, Olivia M. Bertolla, Norihiko Takemoto, Stephan Brouwer, Jack O’Donohue, Masayuki Ono, Eri Ikeda, Victor Nizet, Mark J. Walker, Shigetada Kawabata

## Abstract

*Streptococcus pyogenes* (group A *Streptococcus*; GAS) requires flexible metabolic regulation to adapt to diverse host environments; however, its physiological responses to lactose and glycerol remain poorly characterized compared with glucose. In this study, we analyzed three representative serotype M1 GAS strains—the M1_global_ strain 5448, the ancestral M1 strain SF370, and SP1380 as a representative of the recently emerged M1_UK_ lineage. Glycerol consistently suppressed lactose-dependent growth across all strains, while glycerol-dependent respiratory activity was observed in 5448 and SF370 but was absent in SP1380. RNA-seq analysis of strain 5448 revealed no significant transcriptional changes upon glycerol supplementation, indicating that this inhibitory effect likely occurs through a non-transcriptional mechanism. In contrast, lactose supplementation induced distinct transcriptional programs compared with glucose, including coordinated expression changes gene sets regulated by carbohydrate-responsive transcription factors (CcpA, MalR, and NanR) and by the virulence regulator Mga and the stress-response regulator Rgg. Together, these findings identify a previously unrecognized layer of carbon source-dependent metabolic regulation and growth control in GAS.

## Introduction

GAS is a β-hemolytic Gram-positive bacterium that commonly colonizes the human nasopharynx and skin, particularly in children^1–3^ 【PMID: 27312939, 37246259】. Although often asymptomatic, GAS is a major human pathogen responsible for diseases ranging from mild pharyngitis and impetigo to life-threatening conditions such as necrotizing fasciitis and streptococcal toxic shock syndrome^4^ 【 PMID: 24696436 】. The M1 serotype is a predominant cause of invasive infections globally and in high-income countries, making it essential to understand how this lineage adapts to various host niches to inform new approaches for effective control.

As a bacterium capable of both commensalism and invasive disease, GAS must deploy flexible metabolic strategies to survive in nutrient-diverse environments. Carbohydrate utilization is central to GAS colonization and proliferation^5^ 【PMID: 34463991】, and ∼15% of its core genome encodes proteins involved in carbon metabolism^6^ 【PMID: 16861648 】. GAS preferentially metabolizes glucose through glycolysis^7,8^ 【 PMID: 29594067, 37278526】; however, glucose concentrations in the skin^9,10^ 【PMID: 14598873, 33789094】 and nasopharynx^11^ 【PMID: 14678009】—its initial colonization sites—are markedly lower than in blood, resulting in limited glucose availability for bacterial metabolism. In the human pharynx^1,2^ 【 PMID: 27312939, 37246259 】, GAS also intermittently encounters lactose derived from breast milk or dairy products, and glycerol is detectable in the oral cavity at low millimolar levels and is considered abundant^12^ 【PMID: 39171915】. Despite this, the growth behavior and transcriptional responses of GAS to lactose and glycerol remain largely uncharacterized.

Recent epidemiological studies have reported the global emergence and dissemination of a novel GAS M1 lineage, designated M1_UK_^13–16^ 【PMID: 38729927, 37640370, 36828918, 31519541】. This lineage has rapidly expanded in multiple countries, including the United Kingdom, the United States, and Australia, and is associated with increased incidence of scarlet fever and invasive streptococcal infections^17–19^ 【PMID: 37019153, 38440985, 38035130 】. The M1_UK_ GAS lineage has been suggested to exhibit altered metabolic characteristics compared with the classical M1_global_ lineage. Genomic analyses have identified 27 lineage-defining single-nucleotide polymorphisms (SNPs) in M1_UK_, among which a defining feature is the increased expression of the streptococcal pyrogenic exotoxin A (SpeA)^20^ 【PMID: 37093716 】. Notably, several SNPs occur in genes potentially involved in metabolism, including a nonsense mutation in *gldA* that introduces a premature stop codon and is predicted to disrupt glycerol dehydrogenase function^20^ 【 PMID: 37093716】. In addition, a SNP within the promoter region of *glpF.2* (also known as *glA*), predicted to encode an aquaporin, has been shown to reduce the expression of this gene^15^【PMID: 36828918】. Together, these findings strongly suggest that glycerol metabolism has been remodeled in the M1_UK_ lineage. However, the physiological consequences of these metabolic differences, and how they impact the pathogenicity and adaptability of M1_UK_, remain poorly understood.

Based on these considerations, the present study aimed to elucidate how lactose and glycerol influence the growth and metabolic regulation of GAS M1 serotype strains.

## Results

### Glycerol impairs lactose-dependent growth in all three M1 strains

To investigate how glycerol affects GAS growth on different carbohydrates, the M1_global_ type strain 5448, the ancestral strain SF370, and the newly emergent M1_UK_ clinical isolate SP1380 were cultured in chemically defined medium (CDM)^21,22^ 【PMID: 39158303, 30667377 】 supplemented with glucose, lactose, or glycerol, either individually or in combination (**Fig. 1A**). In CDM lacking a carbon source (Control) or supplemented with glycerol alone, no increase in optical density (OD₆₀₀) was observed, indicating that glycerol cannot support growth under these conditions (**Fig. 1B–D**). Although glycerol had no measurable effect on glucose-dependent growth, it significantly suppressed lactose-dependent growth—a phenotype consistently observed across all three strains (**Fig. 1B–D**).

**Figure 1.**
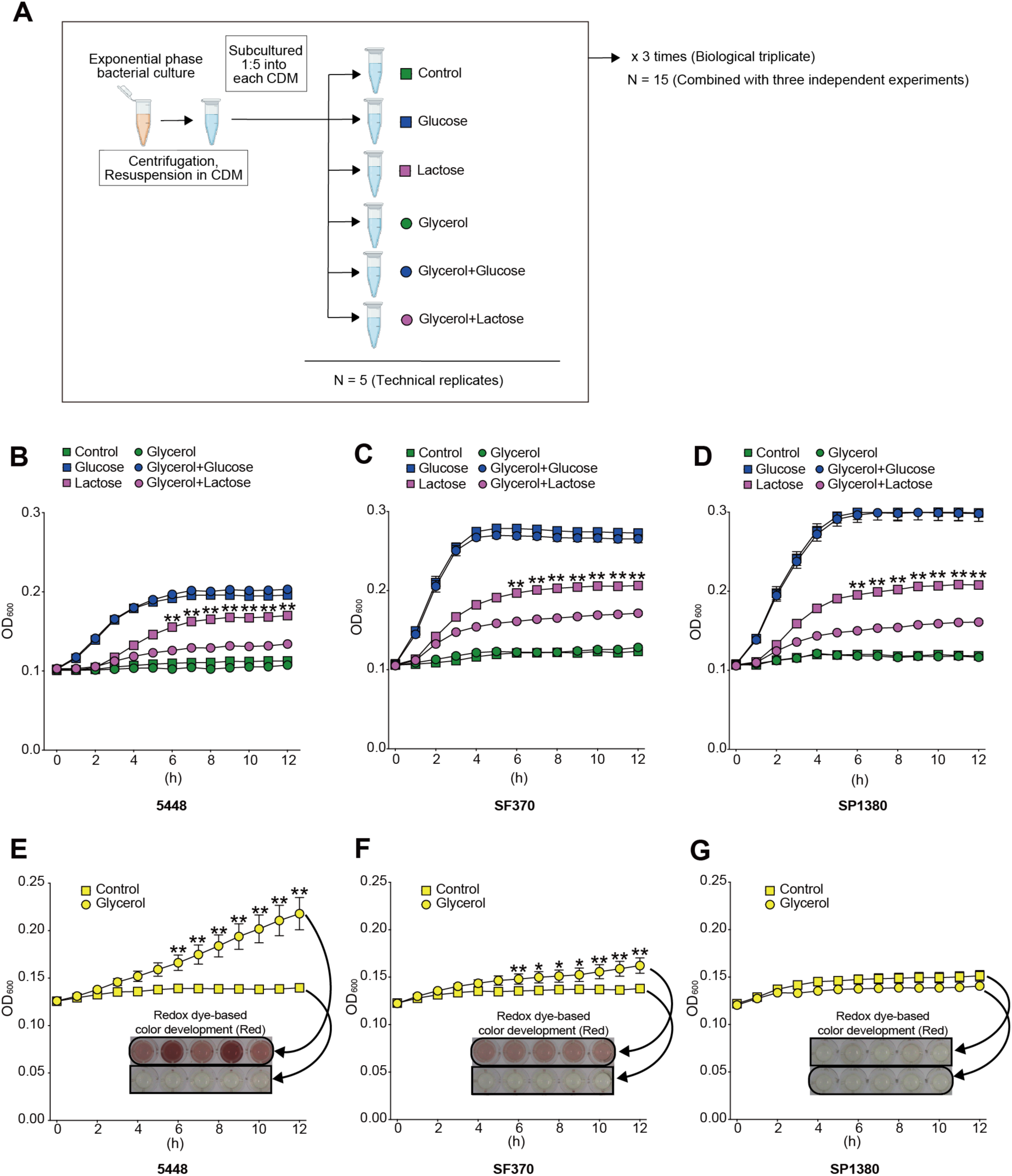
Growth of GAS in different carbohydrate conditions and glycerol-induced respiratory activity (A) Experimental workflow of the growth assay. Exponentially growing cultures were centrifuged, resuspended in chemically defined medium (CDM), and subcultured into CDM supplemented with glucose, lactose, or glycerol, either individually or in combination. Growth curves were monitored under each condition. (B-D) Growth curves of the M1 strain 5448 (B), the ancestral strain SF370 (C), and the M1_UK_ strain SP1380 (D). Glycerol alone did not support growth, whereas glucose supported robust growth. Glycerol had no measurable effect on glucose-dependent growth but significantly impaired lactose-dependent growth in all three strains. (E-G) Respiratory activity assays for each strain. Cultures were incubated in CDM supplemented with Redox Dye H with or without glycerol, and OD₆₀₀ was monitored using a plate reader. Glycerol induced a marked increase in respiratory activity in strains 5448 (E) and SF370 (F), but not in SP1380 (G). Representative images of redox dye-based color development are shown below each graph. Experiments were performed in three independent biological replicates, each with five technical replicates (total n = 15). Error bars indicate the mean ± SEM. Statistical comparisons were performed for time points after 6 h. Statistical differences between groups were analyzed using the Mann-Whitney’s *U*-test. \*\**p* < 0.01, \**p* < 0.05.

### Glycerol induces respiration in 5448 and SF370 but not in SP1380

Previous work from our group has demonstrated that strain 5448 can utilize glycerol as a respiratory substrate in the presence of Redox Dye H^21^ 【PMID: 39158303】. Although a different instrument and plate format were used in the present study, strain 5448 showed similar behavior (**Fig. 1E**), exhibiting a time-dependent increase in red color intensity, indicative of enhanced respiratory activity. Strain SF370 also actively respired glycerol in CDM supplemented with Redox Dye H (**Fig. 1F**). In contrast, SP1380 displayed no change in color intensity under the same conditions, suggesting that glycerol does not trigger respiratory activity in this M1_UK_ lineage strain (**Fig. 1G**).

These findings suggest that, compared with 5448 and SF370, strain SP1380 may have a defect in glycerol uptake or subsequent metabolic processing, consistent with prior genomic reports identifying a nonsense mutation in *gldA* and a promoter SNP in *glpF.2* (*glA*) that are predicted to impair glycerol metabolism^15,20^ 【PMID: 36828918, 37093716】.

### Glycerol utilization pathways in M1 strains of GAS and their regulatory disruption in strain SP1380

According to NCBI sequence annotations, the major enzymes predicted to contribute to glycerol metabolism in GAS M1 strains are glycerol dehydrogenase (*gldA*), glycerol kinase (*glpK*), α-glycerophosphate oxidase (*glpO*), and the glycerol transporters *glpF* and *glpF.2* (**Fig. 2A**). To predict how these strains metabolize glycerol, we used our previously reported genome-scale metabolic model of M1 GAS^21^ 【PMID: 39158303】 together with the Escher visualization platform^23^ 【PMID: 26313928】 to map metabolic routes from glucose, lactose, and glycerol to central metabolism (**Fig. 2B**). The M1 GAS lineage (including strains SF370, 5448, and SP1380) encodes two aquaporin-type glycerol transporters, GlpF and GlpF.2, the latter hypothesized to contribute to glycerol uptake. In the M1_UK_ lineage, a promoter SNP that reduces *glpF.2* (*glA*) expression and a nonsense mutation in *gldA.* Both mutations are present in SP1380^15,20^ 【PMID: 36828918, 37093716】.

**Figure 2.**
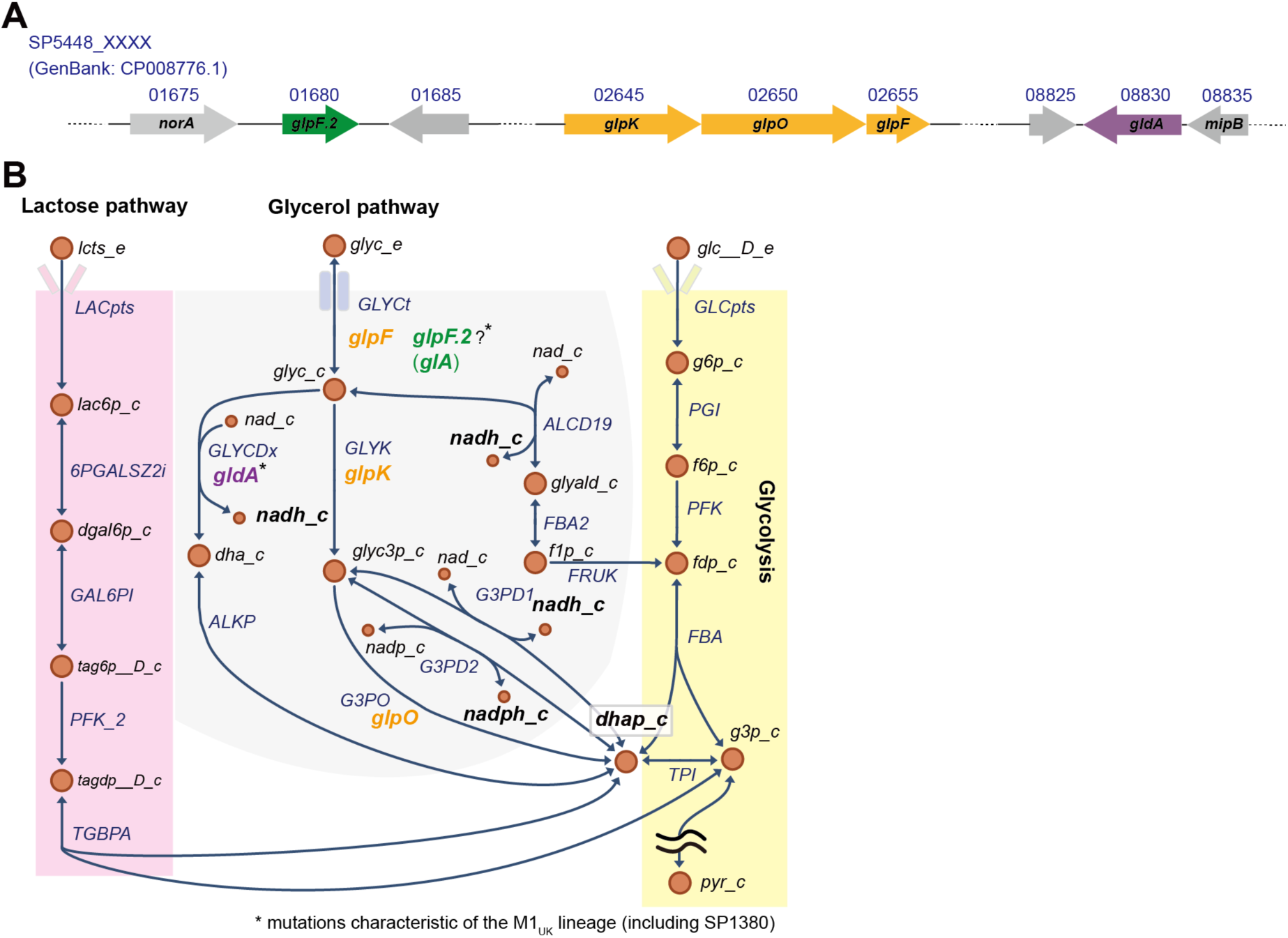
Genetic organization and metabolic overview of carbohydrate utilization in GAS serotype M1. (A) Genomic locus of genes involved in glycerol metabolism. (B) Overview of lactose, glucose, and glycerol metabolism in GAS serotype M1. Metabolite names are annotated with the suffix “_e” for extracellular and “_c” for cytoplasmic localization. Lactose pathway (pink background): Lactose (lcts_e) is transported into the cell via LACpts (PEP:pyruvate phosphotransferase system), forming lactose 6-phosphate (lac6p_c). This intermediate is hydrolyzed by 6PGALSZ2i (6-phospho-β-galactosidase) to produce 6-phospho-D-galactose (dgal6p_c), which is subsequently isomerized by GAL6PI (galactose-6-phosphate isomerase) to yield D-tagatose 6-phosphate (tag6p D_c). PFK_2 (tagatose-6-phosphate kinase) phosphorylates this compound to generate D-tagatose 1,6-bisphosphate (tagdp D_c), which is then cleaved by TGBPA (tagatose-bisphosphate aldolase) into glyceraldehyde 3-phosphate (glyc3p_c) and dihydroxyacetone phosphate (dhap_c), feeding into central metabolism. Glycerol pathway (gray background): Glycerol (glyc_e) is taken up into the cytoplasm via GLYCt encoded by *glpF*, resulting in cytoplasmic glycerol (glyc_c). In addition, *glpF.2* (*glA*) (aquaporin) has been suggested to contribute to glycerol transport, although this remains hypothetical. Glycerol can be metabolized through multiple oxidative routes: (i) GLYCDx (glycerol dehydrogenase, encoded by *gldA*) oxidizes glycerol to dihydroxyacetone (dha_c), which can then be converted to dhap_c by ALKP (alkaline phosphatase) as predicted in our model. (ii) GLYK (glycerol kinase, encoded by *glpK*) first phosphorylates glycerol to produce glycerol 3-phosphate (glyc3p_c), which is then oxidized to dhap_c by G3PD1 or G3PD2 (glycerol-3-phosphate dehydrogenases). (iii) GLYK also phosphorylates glycerol to glyc3p_c, which can alternatively be oxidized by G3PO (glycerol 3-phosphate oxidase, encoded by *glpO*) to generate dhap_c. (iv) ALCD19 (alcohol dehydrogenase) oxidizes glycerol to glyceraldehyde (glyald_c), which is further processed by FBA2 and FRUK to produce fructose 1,6-bisphosphate (fdp_c) via fructose 1-phosphate (f1p_c). Genes related to glycerol metabolism are labeled in green, orange, and purple: *glpF.2* (*glA*) in green, *glpK*, *glpO*, and *glpF* in orange, and *gldA* in purple. Asterisks indicate mutations characteristic of the M1_UK_ lineage, including SP1380: a nonsense mutation in *gldA* and reduced expression of *glpF.2* due to a promoter SNP have been reported^15^ 【PMID: 36828918】. Glucose pathway (yellow background): D-glucose (glc D_e) is taken up via GLCpts (glucose transport via PEP:Pyr PTS), phosphorylated to D-glucose 6-phosphate (g6p_c) by PGI (glucose-6-phosphate isomerase), then to D-fructose 1,6-bisphosphate (fdp_c) by PFK (phosphofructokinase). This is cleaved by FBA (fructose-bisphosphate aldolase) into g3p_c and dhap_c. TPI (triose-phosphate isomerase) interconverts g3p_c and dhap_c, which are then funneled into downstream energy metabolism, ultimately leading to the formation of pyruvate (pyr_c). Arrows indicate the direction of enzymatic reactions and metabolic flux.

Based on information from the genome-scale model, glycerol can enter central metabolism through three potential routes In the first route, glycerol is sequentially processed by ALCD19 (alcohol dehydrogenase), FBA2 (D-fructose-1-phosphate D-glyceraldehyde-3-phosphate lyase), and FRUK (Fructose-1-phosphate kinase) to generate D-fructose 1,6-bisphosphate (fdp_c), a key glycolytic intermediate. In the second route, glycerol is phosphorylated by GLYK (glycerol kinase, *glpK*) to produce glycerol 3-phosphate (glyc3p_c), which is then oxidized to dihydroxyacetone phosphate (dhap_c) by either G3PO (glycerol 3-phosphate oxidase, *glpO*) or G3PD1/2 (glycerol-3-phosphate dehydrogenases, *gpsA*). In the third pathway, glycerol is oxidized directly to dihydroxyacetone (dha_c) by GLYCDx (glycerol dehydrogenase, *gldA*) and then converted to dhap_c. Because the M1_UK_ lineage strains carry a nonsense mutation in *gldA*, this branch is predicted to be impaired^20^【PMID: 37093716】.

Collectively, these findings suggest that although M1 GAS strains possess multiple glycerol utilization pathways, their function may be disrupted in the M1_UK_ lineage. In SP1380, the absence of glycerol-induced respiration (**Fig. 1G**), together with loss of *gldA* function indicates that all three predicted glycerol metabolic routes may be non-functional.

### Glycerol does not influence the GAS transcriptional program

To evaluate transcriptional changes associated with glycerol uptake, we selected strain 5448, which exhibited the most pronounced glycerol-dependent respiratory activity, for RNA-seq analyses. To capture the early transcriptional responses to different carbon sources, RNA was extracted 2 h after inoculation (**Fig. 3A and 3B**), a time point chosen to sensitively reflect carbon source-dependent metabolic adaptation. In addition, because glycerol-dependent inhibition of lactose-supported growth became apparent later in culture, RNA was also collected at 5 h after inoculation for transcriptomic profiling (**Fig. 3D and 3E**).

**Figure 3.**
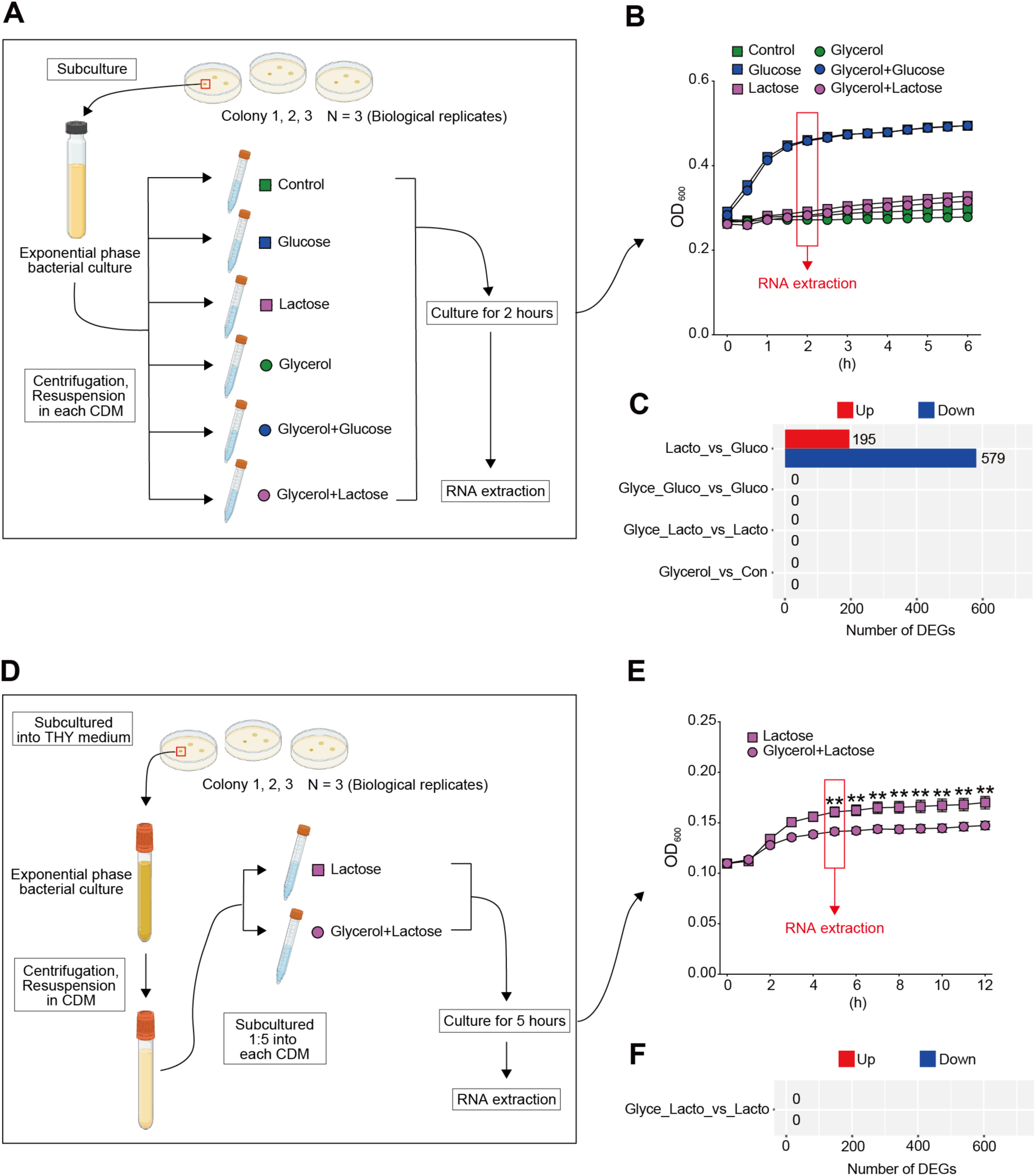
Transcriptomic profiling of GAS strain 5448 under defined carbon source conditions at 2 h and 5 h post-incubation. (A, B) Experimental design for transcriptomic analysis at 2 h. (A) Workflow of RNA-seq experiments. Exponentially growing cultures of GAS 5448 were subcultured into chemically defined medium (CDM) supplemented with glucose, lactose, glycerol, or their combinations (glycerol + glucose, glycerol + lactose). After 2 h of cultivation, cells were harvested for RNA extraction. Three independent biological replicates were performed. (B) Growth curve of the 5448 strain under the indicated conditions. The red arrow indicates the sampling point (2 h). At this time, cultures grown in glucose were in the late-exponential phase, whereas cultures grown in other carbon sources (lactose, glycerol, or glycerol + lactose) remained in the lag phase. RNA was extracted at this point for transcriptomic analysis. Error bars indicate the mean ± SEM (n = 15). (C) Number of differentially expressed genes (DEGs) identified in each pairwise comparison (adjusted *p* < 0.1, |log₂FC| > 1). (D, E) Experimental design for transcriptomic analysis at 5 h. (D) Workflow of RNA-seq experiments. Exponentially growing cultures of GAS 5448 were centrifuged, the supernatant was removed, and cells were resuspended in CDM. Cultures were then subcultured 1:5 into CDM supplemented with lactose or glycerol + lactose. After 5 h of cultivation, cells were harvested for RNA extraction. Three independent biological replicates were performed. (E) Growth curve of the 5448 strain under the indicated conditions. The red arrow indicates the sampling point (5 h). At this time, lactose-dependent growth was suppressed by glycerol, and cells were harvested for RNA extraction for transcriptomic analysis. Error bars indicate the mean ± SEM (n = 15). Statistical comparisons were performed for time points after 5 h. Statistical differences between groups were analyzed using the Mann-Whitney’s *U*-test. \*\**p* < 0.01. (F) DEGs identified between lactose and glycerol + lactose conditions (adjusted *p* < 0.1, |log₂FC| > 1).

At 2 h, no differentially expressed genes (DEGs) were identified when comparing glucose alone with glycerol + glucose, or lactose alone with glycerol + lactose (**Fig. 3C**). Likewise, at 5 h, no DEGs were detected between lactose-supplemented and glycerol + lactose-supplemented conditions (**Fig. 3F**). These findings indicate that glycerol does not measurably alter the global transcriptional profile, even at the time point when its inhibitor effect on glycerol-dependent growth becomes evident.

In contrast, numerous DEGs were detected at 2 h when comparing glucose- and lactose-supplemented conditions, demonstrating the induction of distinct transcriptional programs under these carbon sources (**Fig. 3C**). Principal component analysis (PCA) and hierarchical clustering further revealed clear separation of samples grown in glucose vs. lactose (**Supplementary Fig. 1**), confirming that global transcriptional profiles differ markedly between these conditions.

### Transcriptional changes and iModulon activation in lactose-supplemented cultures compared to glucose

Substantial transcriptional differences were observed between glucose- and lactose-supplemented cultures, with 195 genes significantly upregulated and 579 genes downregulated under lactose conditions (**Fig. 3C**). A full list of significant DEGs is provided in Supplementary Data 1. Exploring these differences may offer insights into why glycerol suppresses lactose-dependent growth.

To investigate regulatory drivers of these transcriptional shifts, we examined iModulons—sets of coexpressed genes identified through independent component analysis (ICA) and linked to specific regulators or biological processes^8,24–26^ 【PMID: 31797920, 32616573, 33311500, 37278526 】. In this framework, iModulon activity is calculated relative to the no-carbon Control (activity = 0), allowing direct comparison across conditions. Using Differential iModulon Activity (DiMA) analysis, we compared iModulon activation under glucose-vs. lactose-supplemented conditions (**Fig. 4A**). Multiple iModulons associated with LacI family transcription factors—including CcpA, MalR, and NanR—showed increased activity under lactose conditions, consistent with the known roles of these regulators in responding to alternative sugars in the *Streptococcus* genus^27–29^ 【PMID: 26030923, 24642967, 25724955 】. Additionally, increased activity was observed in iModulons associated with the virulence regulator Mga (involved in host adhesion and immune evasion)^30^ 【PMID: 9260962】 and Rgg (involved in stress response)^31^ 【PMID: 23188510】. These results suggest that lactose availability triggers a broader coordinated regulatory response.

**Figure 4.**
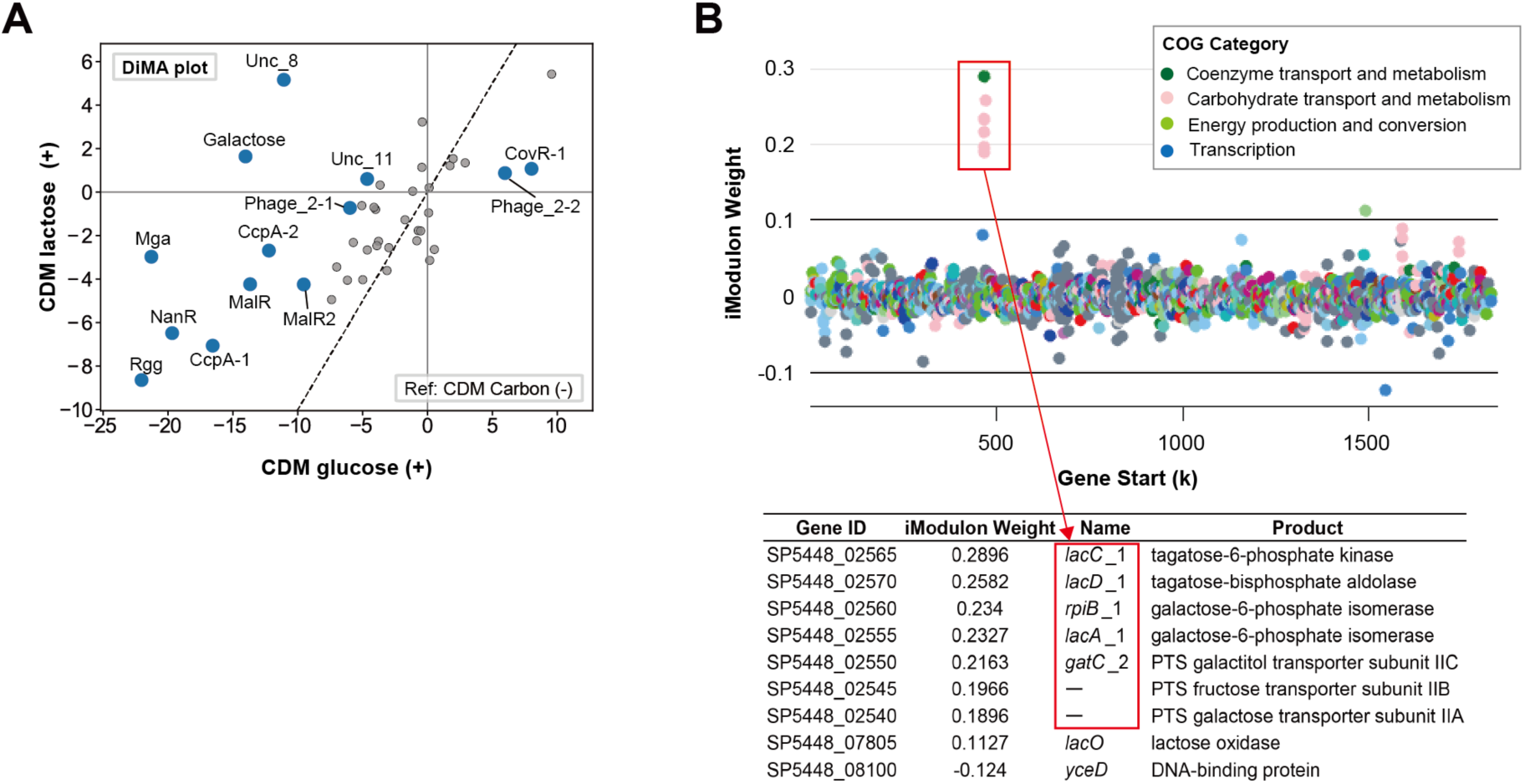
Differential iModulon activity (DiMA) analysis of GAS strain 5448 comparing lactose-supplemented and glucose-supplemented conditions, and genes comprising the Galactose iModulon. (A) The DiMA plot shows the comparison of iModulon activities between lactose- and glucose-grown cultures, based on RNA-seq data from three biological replicates per condition. iModulons associated with carbohydrate-responsive transcription factors (CcpA, MalR, NanR) and host-response regulators (Mga, Rgg) are activated. Notably, the Galactose iModulon also exhibits significant activation under lactose conditions. (B) Distribution and list of genes within the Galactose iModulon, including those involved in galactose and tagatose metabolism.^3^ 【PMID: 39494532】

Galactose iModulon activity was also elevated under lactose conditions. Since lactose is metabolized intracellularly to lactose-6-phosphate, then to galactose-6-phosphate before entering the tagatose-6-phosphate pathway (**Fig. 2B**), induction of galactose-associated metabolic genes is expected and consistent with the observed increase in Galactose iModulon activity. Consistent with this, the galactose-associated metabolic genes *lacA, lacC,* and *lacD* were included in the Galactose iModulon (**Fig. 4B**)^3^ 【PMID: 39494532】.

Together, these findings demonstrate that shifting the available carbon source from glucose to lactose elicits extensive reprogramming of carbohydrate metabolism and regulatory networks, allowing the bacterium to maintain metabolic homeostasis under altered nutrient conditions.

## Discussion

In this study, we investigated how lactose and glycerol influence growth and metabolic regulation in GAS. Among the three representative M1 strains (5448, SF370, and SP1380), glycerol consistently suppressed lactose-dependent growth. Glycerol-dependent respiratory activity was detectable in 5448 and SF370 but absent in SP1380. RNA-seq analysis of 5448 revealed no significant transcriptional changes following glycerol supplementation, suggesting that this inhibitory effect occurs through a non-transcriptional mechanism. Transcriptomic profiling further showed that, compared with glucose-supplemented conditions, lactose-supplemented conditions induced distinct expression changes in multiple coexpressed gene sets (iModulons). These included modules associated with carbohydrate-responsive regulators such as CcpA, MalR, and NanR^27–29^ 【PMID: 26030923, 24642967, 25724955 】, as well as iModulons linked to the virulence regulator Mga^30^ 【PMID: 9260962】 and the stress-response regulator Rgg^31^ 【PMID: 23188510】. Collectively, these findings suggest that, in the absence of glucose, lactose availability induces a coordinated transcriptional program that may enhance host adaptation and influence pathogenic potential.

In the M1_UK_ lineage, including SP1380 strain, a nonsense mutation in *gldA* and reduced expression of *glpF.2* (*glA*) have been reported^15,20^ 【PMID: 37093716, 36828918】. Given the absence of glycerol-dependent respiration in SP1380 under our experimental conditions, it is possible that only the *gldA*-mediated pathway remains functional among the three predicted routes for glycerol metabolism (**Fig. 2B**). However, confirming this hypothesis will require additional studies using *gldA* deletion mutants in 5448 and SF370 backgrounds, which would allow direct comparison of glycerol utilization across the M1_global_, M1ancestral, and M1_UK_ lineages.

As revealed by the DiMA analysis, activation of the Galactose iModulon was also observed under lactose-supplemented conditions. According to iModulonDB (https://imodulondb.org/)^3^ 【PMID: 39494532】, this iModulon includes *lacA*, *lacC*, and *lacD*, components of the *lacABCD* operon, which converts lactose-derived intermediates—galactose-6-phosphate and tagatose-6-phosphate—into glycolytic intermediates. This operon is strongly induced under lactose-utilizing conditions and is conserved across many Gram-positive bacteria, including *Lactococcus lactis* and *Staphylococcus aureus*, where it contributes to lactose catabolism^32,33^ 【PMID: 1901863, 1655695】. In GAS, two distinct *lacABCD*-containing loci exist, referred to as *lac.1* and *lac.2* operons, which have been reported to contribute to lactose-dependent growth^34^ 【 PMID: 17371500 】. These observations are consistent with the metabolic map in Fig. 2B and support the model in which GAS uses the galactose-tagatose pathway to convert lactose into intermediates feeding into glycolysis. Interestingly, the *lac.1* genes were included in the CcpA-2 and Galactose iModulons, whereas the *lac.2* genes were included in the Mga iModulon. All three iModulons were activated under lactose conditions, suggesting roles for CcpA and Mga in regulating lactose-responsive genes. This is consistent with prior work showing that CcpA coordinates virulence and metabolic gene expression in response to carbohydrate availability in GAS^35^【PMID: 19841076】. Moreover, the concurrent activation of CcpA- and Mga-associated iModulons suggests that CcpA may indirectly influence Mga activity through metabolic signaling, likely by altering intracellular carbohydrate flux or energy status. This supports the model in which CcpA integrates metabolic control with virulence gene regulation^35^【PMID: 19841076】. Together, these findings support a model in which CcpA- and Mga-mediated regulatory networks cooperatively control transcriptional responses to lactose in GAS.

From a metabolic perspective, although lactose metabolism also provides some glucose-derived carbon that enters glycolysis upstream, bacterial growth under lactose conditions does not reach the level achieved with glucose alone. Lactose, a disaccharide of glucose and galactose, is metabolized in GAS into glucose-6-phosphate (G6P) and galactose-6-phosphate (Gal6P), respectively. G6P can be directly funneled into glycolysis, supporting efficient ATP production. In contrast, Gal6P enters the tagatose-6-phosphate pathway and is converted into dihydroxyacetone phosphate (DHAP) and glyceraldehyde-3-phosphate (G3P), which merge into glycolysis at downstream steps. Considering this pathway, lactose metabolism should, in principle, be sufficient to support bacterial growth. However, our RNA-seq analysis revealed that the expression of several genes encoding upper glycolytic enzymes—including *pgi* (glucose-6-phosphate isomerase), *pfkA* (6-phosphofructokinase), and *fba* (fructose-bisphosphate aldolase class II)—was decreased under lactose-supplemented conditions compared with glucose-supplemented conditions (**Supplementary Data 2**). These enzymes are responsible for maintaining the levels of glycolytic intermediates such as G6P and fructose-6-phosphate (F6P), which are essential for both energy generation and cell wall biosynthesis. Therefore, impaired synthesis of these intermediates may underlie the reduced growth observed under lactose-supplemented conditions. The underlying factors may not be fully explained by pathway architecture alone and will require further investigation.

Overall, this study demonstrated that glycerol specifically suppresses lactose-dependent growth in GAS through a mechanism that appears independent of transcriptional changes. While we predicted carbon source-specific metabolic pathways and examined the associated metabolic and regulatory responses under different sugar conditions, the precise target by which glycerol inhibits lactose-supported growth could not be identified. Nevertheless, the marked transcriptional reprogramming observed when the carbon source shifted from glucose to lactose indicates that GAS activates coordinated regulatory programs and flexibly modulates its metabolism according to carbohydrate availability. These findings reveal a previously unrecognized aspect of metabolic control in this pathogen and provide an important foundation for deeper investigation into the nutrient-dependent infection strategies of GAS.

## Materials and Methods

### Strains

GAS serotype M1 strain 5448 is widely recognized as part of the modern globally disseminated M1 lineage, commonly termed the M1_global_ or M1T1 clone (accession no. CP008776)^36^ 【PMID: 11035746】. By contrast, GAS M1 strain SF370 is regarded as an ancestral form of the M1 serotype. In the literature SF370 is often described as an “archaic” or pre-epidemic M1 lineage representative^37^ 【PMID: 11296296】. GAS serotype M1 strain SP1380 is a representative of the recently emerged M1_UK_ lineage, which is genetically distinct from the classical M1_global_ clone (accession no. NZ_CP060269)^15^ 【 PMID: 36828918】. Bacterial colonies grown on Tryptic Soy Agar supplemented with 5% sheep blood were inoculated into Todd–Hewitt broth supplemented with 2% yeast extract (THY; Hardy Diagnostics, Santa Maria, CA, USA) and cultured overnight at 37°C in screw-cap glass tubes under static conditions. The overnight cultures were used for subsequent experiments.

### Growth assay

Growth experiments were performed in a modified chemically defined medium (CDM), based on the formulation reported by Zhu et al.^22^ 【PMID: 30667377】, with all component concentrations reduced to one-fourth as previously described^21^ 【PMID: 39158303】. CDM was supplemented with 4.5 g/L of glucose, lactose, or glycerol as sole or combined carbon sources. GAS cultures grown overnight in THY were diluted 1:5 into fresh THY and incubated at 37°C until they reached the mid-exponential phase (OD₆₀₀ = 0.4). Cells were harvested by centrifugation and resuspended in CDM without any carbon source. The resulting suspension was then diluted 1:5 into CDM containing the designated carbon sources.

Growth kinetics were assessed in 96-well microplates using a Tecan Infinite Pro 200 plate reader (Tecan, Männedorf, Switzerland). Plates were sealed and incubated at 37°C, with OD₆₀₀ measured at 30-minute intervals.

### Respiration assay

To evaluate bacterial respiratory activity, we used Biolog Redox Dye H (Biolog Inc., Hayward, CA, USA), which detects cellular respiration through colorimetric reduction. GAS cultures were grown overnight at 37°C in THY, and the overnight cultures were harvested by centrifugation and resuspended in CDM without any carbon source. The resulting suspension was then diluted 1:10 into CDM mixed with Redox Dye H at a final concentration of 1% (v/v). Glycerol was supplemented with 4.5 g/L in CDM. Respiration kinetics were monitored using a Tecan Infinite Pro 200 plate reader (Tecan, Männedorf, Switzerland) at 37°C, with OD₆₀₀ measurements taken every 30 minutes. Microplates were sealed during incubation to prevent evaporation and maintain sterility.

### RNA extraction and library preparation

Overnight cultures in THY medium derived from different colonies were diluted 1:5 into fresh THY medium and incubated at 37°C, and growth was monitored by measuring the optical density at 600 nm. Mid-exponential phase cells (OD₆₀₀ = 0.4) were harvested by centrifugation, resuspended in CDM supplemented with the indicated carbohydrate. For RNA extraction at 2 h, cultures were directly transferred to each carbohydrate condition at 1× concentration to obtain sufficient biomass. For RNA extraction at 5 h, cultures were inoculated at a 1:5 dilution and incubated under the same conditions.

Each sample was then mixed with twice its volume of RNAprotect Bacteria Reagent (Qiagen, Hilden, Germany), vortexed, and incubated at room temperature for 5 min. After centrifugation, the supernatant was discarded, and total RNA was extracted from the resulting cell pellets using the Quick RNA Fungal/Bacterial Microprep kit (Zymo Research, Irvine, CA, USA). Cells were disrupted mechanically for 30 s using a Multi Beads Shocker® MB3000 (Yasui Kikai, Osaka, Japan). Library preparation was performed with the Ribo-Zero Plus Microbiome kit and the TruSeq Stranded mRNA Library Prep kit (Illumina, San Diego, CA, USA) according to the manufacturers’ protocols. Sample details of the 24 RNA samples and DEG information are provided in Supplementary Data 1. All samples were collected in biological triplicates.

### RNA-seq and data analysis

RNA sequencing and primary data analysis were performed at the NGS core facility of the Research Institute for Microbial Diseases, Osaka University. Libraries were prepared using the RiboZero Plus Microbiome kit followed by TruSeq Stranded mRNA workflow (Illumina) and sequenced on the NovaSeq X Plus platform (Illumina) using 151 bp paired-end reads. Data processing and quality filtering were performed based on a previously published pipeline^38^ 【PMID: 31028276】, with slight adaptations. Raw sequence reads were quality-trimmed using fastp, and then mapped to the GAS strain 5448 genome using Bowtie2^39^ 【PMID: 22388286】. Read counts were assigned to open reading frames (ORFs) using featureCounts^40^ 【PMID: 24227677】. Genes with an average count below 10 across samples were excluded to reduce noise from low-expression features. Transcript abundance was normalized and expressed in transcripts per million (TPM). Differential gene expression and global transcriptomic analyses were conducted using iDEP^41^ 【PMID: 33835455】. For dimensionality reduction and clustering, EdgeR log transformation was applied, and hierarchical clustering was performed using the average linkage method with correlation distance.

RNA-seq data generated in this study have been deposited in the DDBJ Sequence Read Archive (DRA) under BioProject accession number PRJDB37997, and individual sequencing runs are available under accession numbers DRR793594–DRR793617. Data will be released on 31 October 2027 in accordance with DDBJ hold policy.

### Data for iModulon Analysis and Code Availability

We utilized statistically independent components (M) previously identified through independent component analysis (ICA), along with normalized log TPM values (X) and their associated activity profiles (A) across various experimental conditions, all of which were obtained from iModulonDB^3,8^ 【PMID: 39494532, 37278526】.

In this study, we further incorporated newly obtained normalized log TPM values (X) and the corresponding activity profiles (A), which were calculated using count data derived from bacterial RNA samples collected after 2 h of incubation. Details of these combined datasets are provided in Supplementary Data 2. The code for assessment of the regulator enrichment can be found on Github (https://github.com/avsastry/modulome-workflow). Python package for analyzing and visualizing iModulons (PyModulon) can be also found on Github, (https://github.com/SBRG/pymodulon).

### Quantification and statistical analysis

Statistical analysis was performed using GraphPad Prism version 10.4.1. (GraphPad Software Inc., La Jolla, CA, USA). Differences between groups were analyzed using a Mann-Whitney *U* test. Sample sizes and *p* values are indicated in figure legends.

### Declaration of Generative AI and AI-Assisted Technologies in the Writing Process

During the preparation of this work the author(s) used ChatGPT 5.1 in order to assist with English-language editing. After using this tool/service, the author(s) reviewed and edited the content as needed, and take(s) full responsibility for the content of the publication.

## Acknowledgments

This study was supported in part by the Ministry of Health, Labour and Welfare of Japan and the Japan Agency for Medical Research and Development (AMED) (JP23wm0325066); Japanese Society for the Promotion of Science (JSPS) KAKENHI (25K00123, 25K028040, 23KK0281); Takeda Science Foundation. The funders had no role in study design, data collection or analysis, decision to publish, or preparation of the manuscript. We acknowledge the NGS core facility at the Research Institute for Microbial Diseases of The University of Osaka for the sequencing and data analysis.

## References

1 Brouwer, S., Barnett, T. C., Rivera-Hernandez, T., Rohde, M. & Walker, M. J. *Streptococcus pyogenes* adhesion and colonization. FEBS Lett 590, 3739–3757 (2016). 10.1002/1873-3468.12254

2 Keeley, A. J., et al. *Streptococcus pyogenes* Colonization in Children Aged 24-59 Months in the Gambia: Impact of Live Attenuated Influenza Vaccine and Associated Serological Responses. J Infect Dis 228, 957–965 (2023). 10.1093/infdis/jiad153

3 Catoiu, E. A. et al. iModulonDB 2.0: dynamic tools to facilitate knowledge-mining and user-enabled analyses of curated transcriptomic datasets. Nucleic Acids Res 53, D99–D106 (2025). 10.1093/nar/gkae1009

4 Walker, M. J. et al. Disease manifestations and pathogenic mechanisms of Group A *Streptococcus*. Clin Microbiol Rev 27, 264–301 (2014). 10.1128/CMR.00101-13

5 Sedghi, L., DiMassa, V., Harrington, A., Lynch, S. V. & Kapila, Y. L. The oral microbiome: Role of key organisms and complex networks in oral health and disease. Periodontol 2000 87, 107–131 (2021). 10.1111/prd.12393

6 Shelburne, S. A, 3rd. et al. Maltodextrin utilization plays a key role in the ability of group A Streptococcus to colonize the oropharynx. Infect Immun 74, 4605–4614 (2006). 10.1128/IAI.00477-06

7 Sundar, G. S. et al. Route of Glucose Uptake in the Group a *Streptococcus* Impacts SLS-Mediated Hemolysis and Survival in Human Blood. Front Cell Infect Microbiol 8, 71 (2018). 10.3389/fcimb.2018.00071

8 Hirose, Y. et al. Elucidation of independently modulated genes in *Streptococcus pyogenes* reveals carbon sources that control its expression of hemolytic toxins. mSystems 8, e0024723 (2023). 10.1128/msystems.00247-23

9 Cunningham, D. D. & Young, D. F. Measurements of glucose on the skin surface, in stratum corneum and in transcutaneous extracts: implications for physiological sampling. Clin Chem Lab Med 41, 1224–1228 (2003). 10.1515/CCLM.2003.187

10 Hirose, Y., et al. *Streptococcus pyogenes* upregulates arginine catabolism to exert its pathogenesis on the skin surface. Cell Rep 34, 108924 (2021). 10.1016/j.celrep.2021.108924

11 Wood, D. M., Brennan, A. L., Philips, B. J. & Baker, E. H. Effect of hyperglycaemia on glucose concentration of human nasal secretions. Clin Sci (Lond) 106, 527–533 (2004). 10.1042/CS20030333

12 Taylor, Z. A. et al. Glycerol metabolism contributes to competition by oral streptococci through production of hydrogen peroxide. J Bacteriol 206, e0022724 (2024). 10.1128/jb.00227-24

13 Vieira, A. et al. Rapid expansion and international spread of M1(UK) in the post-pandemic UK upsurge of *Streptococcus pyogenes*. Nat Commun 15, 3916 (2024). 10.1038/s41467-024-47929-7

14 Li, Y. et al. Expansion of Invasive Group A Streptococcus M1(UK) Lineage in Active Bacterial Core Surveillance, United States, 2019‒2021. Emerg Infect Dis 29, 2116–2120 (2023). 10.3201/eid2910.230675

15 Davies, M. R. et al. Detection of *Streptococcus pyogenes* M1(UK) in Australia and characterization of the mutation driving enhanced expression of superantigen SpeA. Nat Commun 14, 1051 (2023). 10.1038/s41467-023-36717-4

16 Lynskey, N. N. et al. Emergence of dominant toxigenic M1T1 *Streptococcus pyogenes* clone during increased scarlet fever activity in England: a population-based molecular epidemiological study. Lancet Infect Dis 19, 1209–1218 (2019). 10.1016/S1473-3099(19)30446-3

17 Zhi, X. et al. Emerging Invasive Group A Streptococcus M1(UK) Lineage Detected by Allele-Specific PCR, England, 2020(1). Emerg Infect Dis 29, 1007–1010 (2023). 10.3201/eid2905.221887

18 Ramirez de Arellano, E., et al. Clinical, microbiological, and molecular characterization of pediatric invasive infections by *Streptococcus pyogenes* in Spain in a context of global outbreak. mSphere 9, e0072923 (2024). 10.1128/msphere.00729-23

19. Lacey, J. A., et al. A worldwide population of Streptococcus pyogenes strains circulating among school-aged children in Auckland, New Zealand: a genomic epidemiology analysis. Lancet Reg Health West Pac 42, 100964 (2024). 10.1016/j.lanwpc.2023.100964

20 Li, H. K. et al. Characterization of emergent toxigenic M1(UK) *Streptococcus pyogenes* and associated sublineages. Microb Genom 9 (2023). 10.1099/mgen.0.000994

21 Hirose, Y. et al. A genome-scale metabolic model of a globally disseminated hyperinvasive M1 strain of *Streptococcus pyogenes*. mSystems 9, e0073624 (2024). 10.1128/msystems.00736-24

22 Zhu, L. et al. Gene fitness landscape of group A *streptococcus* during necrotizing myositis. J Clin Invest 129, 887–901 (2019). 10.1172/JCI124994

23 King, Z. A. et al. Escher: A Web Application for Building, Sharing, and Embedding Data-Rich Visualizations of Biological Pathways. PLoS Comput Biol 11, e1004321 (2015). 10.1371/journal.pcbi.1004321

24 Sastry, A. V. et al. The *Escherichia coli* transcriptome mostly consists of independently regulated modules. Nat Commun 10, 5536 (2019). 10.1038/s41467-019-13483-w

25 Poudel, S. et al. Revealing 29 sets of independently modulated genes in *Staphylococcus aureus*, their regulators, and role in key physiological response. Proc Natl Acad Sci U S A 117, 17228–17239 (2020). 10.1073/pnas.2008413117

26 Rychel, K., Sastry, A. V. & Palsson, B. O. Machine learning uncovers independently regulated modules in the *Bacillus subtilis* transcriptome. Nat Commun 11, 6338 (2020). 10.1038/s41467-020-20153-9

27 Afzal, M., Shafeeq, S., Manzoor, I. & Kuipers, O. P. Maltose-Dependent Transcriptional Regulation of the mal Regulon by MalR in *Streptococcus pneumoniae*. PLoS One 10, e0127579 (2015). 10.1371/journal.pone.0127579

28 Ferrando, M. L. et al. Carbohydrate availability regulates virulence gene expression in *Streptococcus suis*. PLoS One 9, e89334 (2014). 10.1371/journal.pone.0089334

29 Afzal, M., Shafeeq, S., Ahmed, H. & Kuipers, O. P. Sialic acid-mediated gene expression in *Streptococcus pneumoniae* and role of NanR as a transcriptional activator of the nan gene cluster. Appl Environ Microbiol 81, 3121–3131 (2015). 10.1128/AEM.00499-15

30 McIver, K. S. & Scott, J. R. Role of mga in growth phase regulation of virulence genes of the group A *streptococcus*. J Bacteriol 179, 5178–5187 (1997). 10.1128/jb.179.16.5178-5187.1997

31 Lasarre, B., Aggarwal, C. & Federle, M. J. Antagonistic Rgg regulators mediate quorum sensing via competitive DNA binding in *Streptococcus pyogenes*. mBio 3 (2013). 10.1128/mBio.00333-12

32 van Rooijen, R. J., van Schalkwijk, S. & de Vos, W. M. Molecular cloning, characterization, and nucleotide sequence of the tagatose 6-phosphate pathway gene cluster of the lactose operon of *Lactococcus lactis*. J Biol Chem 266, 7176–7181 (1991).

33 Rosey, E. L., Oskouian, B. & Stewart, G. C. Lactose metabolism by *Staphylococcus aureus*: characterization of *lacABCD*, the structural genes of the tagatose 6-phosphate pathway. J Bacteriol 173, 5992–5998 (1991). 10.1128/jb.173.19.5992-5998.1991

34 Loughman, J. A. & Caparon, M. G. Comparative functional analysis of the *lac* operons in *Streptococcus pyogenes*. Mol Microbiol 64, 269–280 (2007). 10.1111/j.1365-2958.2007.05663.x

35 Kietzman, C. C. & Caparon, M. G. CcpA and LacD.1 affect temporal regulation of Streptococcus pyogenes virulence genes. Infect Immun 78, 241–252 (2010). 10.1128/IAI.00746-09

36 Kansal, R. G., McGeer, A., Low, D. E., Norrby-Teglund, A. & Kotb, M. Inverse relation between disease severity and expression of the streptococcal cysteine protease, SpeB, among clonal M1T1 isolates recovered from invasive group A streptococcal infection cases. Infect Immun 68, 6362–6369 (2000). 10.1128/IAI.68.11.6362-6369.2000

37 Ferretti, J. J. et al. Complete genome sequence of an M1 strain of *Streptococcus pyogenes*. Proc Natl Acad Sci U S A 98, 4658–4663 (2001). 10.1073/pnas.071559398

38 Poudel, S. et al. Characterization of CA-MRSA TCH1516 exposed to nafcillin in bacteriological and physiological media. Sci Data 6, 43 (2019). 10.1038/s41597-019-0051-4

39 Langmead, B. & Salzberg, S. L. Fast gapped-read alignment with Bowtie 2. Nat Methods 9, 357–359 (2012). 10.1038/nmeth.1923

40 Liao, Y., Smyth, G. K. & Shi, W. featureCounts: an efficient general purpose program for assigning sequence reads to genomic features. Bioinformatics 30, 923–930 (2014). 10.1093/bioinformatics/btt656

41 Ge, X. iDEP Web Application for RNA-Seq Data Analysis. Methods Mol Biol 2284, 417–443 (2021). 10.1007/978-1-0716-1307-8_22

